# Expansion and differentiation of *ex vivo* cultured erythroblasts in scalable stirred bioreactors

**DOI:** 10.1101/2022.02.11.480112

**Authors:** Joan Sebastián Gallego-Murillo, Giulia Iacono, Luuk A.M. van der Wielen, Emile van den Akker, Marieke von Lindern, Sebastian Aljoscha Wahl

## Abstract

Transfusion of donor-derived red blood cells (RBCs) is the most common form of cell therapy. Production of transfusion-ready cultured RBCs (cRBCs) is a promising replacement for the current fully donor-dependent therapy. However, very large number of cells are required for transfusion. Here we scale-up cRBC production from static cultures to 0.5 L stirred tank bioreactors, and identify the effect of operating conditions on the efficiency of the process. Oxygen requirement of proliferating erythroblasts (0.55-2.01 pg/cell/h) required sparging of air to maintain the dissolved oxygen concentration at the tested setpoint (2.88 mg O_2_/L). Erythroblasts could be cultured at dissolved oxygen concentrations as low as 0.7 O_2_ mg/mL without negative impact on proliferation, viability or differentiation dynamics. Stirring speeds of up to 600 rpm supported erythroblast proliferation, while 1800 rpm led to a transient halt in growth and accelerated differentiation followed by a recovery after 5 days of culture. Erythroblasts could also be differentiated in bioreactors, with final enucleation levels and hemoglobin content similar to parallel cultures under static conditions. After defining optimal mixing and aeration strategies, erythroblast proliferation cultures were successfully scaled up to 3 L bioreactors.

## Introduction

Blood transfusion is the most common cell therapy to this date. Worldwide about 120 million blood donations are collected; nevertheless, availability of blood products for transfusion purposes is not uniformly distributed, with current shortages localized mostly in low-income countries (World Health Organization, 2021). Globalization and an increasing multiethnicity are societal factors that prompt for a larger, more diverse blood supply to ensure a safe transfusion practice (Klinkenberg et al., 2019; Vichinsky et al., 1990). Currently, over 360 blood group antigens are known, being part of more than 35 known blood group systems (Daniels, 2013). Finally, donor-dependent transfusion products harbor a risk for novel bloodborne pathogens that escape current screening programs.

Production of cultured red blood cells (cRBCs) represents a potentially unlimited source of RBCs for transfusion purposes, offering a better control on the quality and safety of the final product. Fully matched cRBCs could be used for patients that have developed alloimmunization or with rare blood groups (Pellegrin et al., 2021; Peyrard et al., 2011). Furthermore, cRBCs loaded with therapeutics or engineered to present antigens in their surface could be used as a highly efficient drug-delivery system (Koleva et al., 2020; Sun et al., 2017; Vichinsky et al., 1990; X. Zhang et al., 2021).

Current *ex vivo* RBC culture protocols start with the commitment of CD34^+^ hematopoietic stem cells (HSCs) to the erythroid lineage, which is characterized by high expression of the transferrin receptor (CD71). This is followed by the proliferation of early erythroblasts expressing glycophorin A (CD235a^dim^) and integrin-α4 (CD49d^dim-to-high^) together with CD71^high^. Finally, the cells mature into CD235^high^/CD49d^-^/CD71^-^ enucleated reticulocytes. Erythropoietin (Epo) is required throughout erythropoiesis for cell survival and proliferation, whereas stem cell factor (SCF) promotes the proliferation capacity (Heshusius et al., 2019; Migliaccio et al., 2002). The addition of glucocorticoids enhances proliferation of early erythroblasts and arrests differentiation. As a result, withdrawal of glucocorticoids and SCF allows for synchronous differentiation (Leberbauer et al., 2005; von Lindern et al., 1999).

Erythroid cells are commonly cultured in static dishes or flasks, in which spatial inhomogeneities in nutrient concentrations occur due to settling of cells to the bottom of the vessels. Moreover, this cultivation system is susceptible to mass transfer limitations as transfer of oxygen to the culture is fully diffusion-dependent (Peniche Silva et al., 2020; Place et al., 2017; Sugiura et al., 2011). This type of culture systems would culminate in impractical numbers of dishes/flasks (currently about 9.000 - 12.000 T175 flasks for a single transfusion unit) (Timmins & Nielsen, 2009, 2011).

The use of a static culture system in which oxygen transfer is facilitated by a gas-permeable membrane at the bottom of the vessel enabled a 50-fold increase in surface cell density (cells per cm^2^) compared to culture dishes (Heshusius et al., 2019). Nevertheless, scale up of this culture protocol is still by area, and limited control on culture conditions can be performed.

Cultivation systems mimicking the microstructure and microenvironment of the bone marrow (BM) niche were developed for cRBC production. Hollow fiber bioreactors enable tissue-like cell concentrations with continuous perfusion of fresh nutrients and dissolved oxygen (Allenby et al., 2019; Housler et al., 2012). Microcarriers and porous scaffolds are used, both under static (Severn et al., 2016, 2019) and agitated conditions (Lee et al., 2015). These biomimetic systems support high erythroblast cell densities. However, scale-up is hampered by gradients in nutrient concentrations along and across hollow fibers (Mohebbi-Kalhori et al., 2012; Piret et al., 1991) or inside microcarriers (Preissmann et al., 1997; Yu, 2012).

Active agitation of RBC cultures increases nutrient homogeneity and oxygen transfer. This includes roller bottles (Y. Zhang et al., 2017), rocking motion bioreactors (Boehm et al., 2010; Timmins et al., 2011), shake flasks (Aglialoro et al., 2021; Sivalingam et al., 2020), and spinner flasks (Griffiths et al., 2012; Kupzig et al., 2017; Sivalingam et al., 2020; Trakarnsanga et al., 2017). Stirred tank reactors (STRs) additionally allow for online monitoring and control of process parameters such as pH, dissolved oxygen and nutrient concentrations. STRs are commonly used for suspension cultures of different mammalian cells such as CHO, HEK293, hybridoma or Vero cell lines (Chu & Robinson, 2001; Rodrigues et al., 2010; Tapia et al., 2016), reaching volumes of up to ∼20.000 L at an industrial scale (Farid, 2007). This type of reactors can be combined with cell retention systems, such as spin filters or external filtration systems, allowing for continuous culture processes with higher cell densities (Avgerinos et al., 1990; Karst et al., 2016).

For cRBCs, STRs were used in small scale cultures (Han et al., 2021; Lee et al., 2018; Ratcliffe et al., 2012, 2012), or in an attempt to scale-up production (Law & Gilbert, 2021). Some studies report a large increase in cell numbers, but the conditions that were used varied widely with respect to medium composition, cell density, oxygenation, agitation (impeller type, speed), and nutrient feeding strategy, making a direct comparison to other cultivation systems challenging (Bayley et al., 2017; Han et al., 2021).

Shear stress is a relevant parameter in scale-up. Hydrodynamic conditions in the BM microenvironment, dominated by slow perfusion of nutrients and low fluid shear stress (0.29 ± 0.27 Pa in sinusoidal capillaries), are different to those of STRs in which complex patterns of fluid flow take place (Bixel et al., 2017; Mazo et al., 1998). The impact of shear stress on cultures seems to be cell-line dependent, affecting cell growth, viability, morphology, protein glycosylation, and differentiation fate (Godoy-Silva et al., 2009; Kretzmer & Schügerl, 1991; Wolfe & Ahsan, 2013; Wu, 1999). Shear stress in rocking culture plates or upon orbital shaking in Erlenmeyer flasks reduced erythroblast viability and accelerated erythroid differentiation (Aglialoro et al., 2021; Boehm et al., 2010). Enhanced differentiation is in agreement with increased enucleation and reduced cell proliferation reported for STRs (Bayley et al., 2017).

The different stages of erythroid cultures show distinct sensitivity to shear stress. Immature erythroid cells responded to shear stress with low viability and increased cell fragility, whereas cultures of more mature cells were robust and showed a linear increase in enucleation through the range of tested stirring speeds (Han et al., 2021). The effect of shear stress on erythroblasts may be mediated, at least partially, by the mechanosensitive channel PIEZO1, which activates multiple calcium-dependent signaling transduction cascades including the Calcineurin-NFAT pathway, the modulation of STAT5 and ERK signaling, and inside-out integrin activation (Aglialoro et al., 2020, 2021; Caulier et al., 2020).

Sparging air also produces shear stress in the cultures, the negative effect of which could be counteracted in cRBC cultures by antifoaming agents at the cost of a lower percentage of enucleated cells (Bayley et al., 2017). The negative effect of turbulence can be aggravated when higher stirring speeds and gas flow rates are needed upon scale-up to keep the cultures well-mixed with adequate oxygen supply (Xing et al., 2009). Nevertheless, it is still unclear if the observed negative effect of sparging on cell cultures is due to the mechanical forces generated during bubble coalescence and breakup, or by biochemical interactions between cells and air bubbles (Sobolewski et al., 2011; Walls et al., 2017; Walsh et al., 2017).

Oxygen availability is a critical parameter in mammalian cell cultures. Cell cultures are often performed under dissolved oxygen (dO_2_) concentrations that are higher than those *in vivo*, potentially leading to an increase in oxidative stress, decrease in growth rate, acceleration of cell differentiation, and increase in apoptosis (Mas-Bargues et al., 2019). *In vivo*, erythroblasts are exposed to the hypoxic conditions of the extravascular bone marrow niche, with a mean oxygen partial pressure of 13.3 mmHg (range: 4.8-21.1 mmHg; (Spencer et al., 2014)), equivalent to a dO_2_ concentration of 0.6 mg O_2_/L (range: 0.2-0.9 mg O_2_/L), or a 8% (range: 3%-13%) of saturation in equilibrium with air (1 atm, 37°C).

Low oxygen pressure in static culture conditions accelerated erythroblast differentiation, increased hemoglobin levels and the enucleation rates, and increased the expression of the hypoxia-inducible factor 1α (HIF1α) (Bapat et al., 2021; Goto et al., 2019). Vlaski et al. also reported accelerated erythroid differentiation, but an increased cell yield in the first culture stage, using gas oxygen concentrations as low as 1.5% (Vlaski et al., 2009). However, it is difficult to predict the local dO_2_ concentrations to which cells are exposed in the culture dishes used for these experiments, as static culture systems often display dO_2_ gradients (Place et al., 2017). In contrast, a more homogeneous dO_2_ concentration can be monitored and controlled in STR cultivations using in-line oxygen measurements. The effect of dO_2_ on enucleation and cell yields in STR erythroid cultures depends on other process parameters such as pH, temperature and shear (Han et al., 2021).

Oxygen requirements of erythroid cultures are dynamic during the maturation process, with cells switching between oxidative phosphorylation (OXPHOS) and glycolysis (Richard et al., 2019). Although oxygen requirements of mature RBCs are well known, there is limited data for erythroid precursors. Oxygen consumption rates for erythroblast have been estimated in Seahorse assays, ranging between 0.26 and 1.66 pg/cell/h (Caielli et al., 2021; Gonzalez-Menendez et al., 2021; Jensen et al., 2019). Erythroblast oxygen requirements in STR cultures range between 0.06 and 0.21 pg/cell/h (cell-specific oxygen consumption rate; q_O2_), and go as low as 0.01-0.05 pg/cell/h in the later stages of differentiation (Bayley et al., 2017).

In the present study, we report the implementation of our cRBC culture protocol in stirred tank reactors. As oxygen availability was identified as a critical parameter controlling the yields of cRBCs, we estimated the oxygen requirements of proliferating erythroblasts, and evaluated the effect of dissolved oxygen and stirring speed on cell yields during erythroblast proliferation and differentiation in this culture system. Based on appropriate culture conditions identified in 0.5 L STRs, we explored the scale up to 3 L STRs, maintaining the same proliferation and differentiation efficiency.

## Materials and methods

### Cell culture

Human adult peripheral blood mononuclear cells (PBMCs) were purified by density centrifugation using Ficoll-Paque (density = 1.077 g/mL; 600g, 30 minutes; GE Healthcare; USA). Informed written consent was given by donors to give approval for the use of waste material for research purposes, and was checked by Sanquin’s NVT Committee (approval file number NVT0258; 2012) in accordance with the Declaration of Helsinki and the Sanquin Ethical Advisory Board. RBCs were cultured from PBMCs as previously described (Heshusius et al., 2019), with minor modifications: nucleosides and trace elements were omitted; cholesterol, oleic acid, and L-α-phosphatidylcholine were replaced by a defined lipid mix (1:1000; Sigma-Aldrich cat#L0288; USA). Expansion cultures were supplemented with erythropoietin (Epo; 2 U/mL; EPREX^®^; Janssen-Cilag; Netherlands), human stem cell factor (hSCF; 50 ng/mL, produced in HEK293T cells), dexamethasone (Dex; 1 µmol/L; Sigma-Aldrich), and interleukin-3 (IL-3; 1 ng/mL, first day only; Stemcell Technologies; Canada). Cell density was maintained between 0.7-2×10^6^ cells/mL by daily feeding with fresh expansion medium.

To induce differentiation, cells were washed and reseeded at 1×10^6^ cells/mL in Cellquin medium supplemented with Epo (10 U/mL), 5% Omniplasma (Octapharma GmbH; Germany), human plasma-derived holotransferrin (1 mg/mL; Sanquin; Netherlands), and heparin (5 U/mL; LEO Pharma A/S; Denmark). Cells were kept in culture for 11 days, without medium refreshment, until fully differentiated, in either stirred bioreactors or in culture dishes.

### Bioreactor and reference culture conditions

All cultivations were performed on either autoclavable (MiniBio 500 mL, or 2 L single wall; glass) or single-use (AppliFlex 0.5 L, or AppliFlex 3.0 L; plastic) stirred bioreactors (Applikon Biotechnology; Netherlands). For expansion cultures, day 8-10 cell cultures from PBMCs were seeded in bioreactors at 0.7-0.9×10^6^ cells/mL. The cell concentration was measured every 24 h, and partial media refreshment was performed if the measured cell density was >1.2×10^6^ cells/mL, diluting the cell culture to 0.7×10^6^ cells/mL. For differentiation cultures, cells were seeded at 1×10^6^ cells/mL and cultured without medium additions.

Process control was established using PIMS Lucullus (Securecell AG; Switzerland). Cultures were agitated using a marine impeller (down-pumping) at defined stirring speeds. pH was measured using an AppliSens pH+ probe (AppliSens; Netherlands) and kept at 7.5 by sparging CO_2_ (acid). pH probe drift was corrected by recalibration every 2 days using an off-line pH measurement. Dissolved oxygen (dO_2_) concentration was monitored using a polarographic probe (AppliSens), and was controlled by sparging pure air using a porous sparger (average pore size = 15 μm). dO_2_ values are reported as the percentage relative to the oxygen saturation concentration in water at equilibrium with air (1 atm, 37°C; 100% = 7.20 mg O_2_/L).

Unless indicated otherwise, 0.5 L STR cultures were performed keeping the working volume constant at 300 mL, with a stirring speed of 200 rpm using a marine impeller of diameter 2.8 cm, and a constant headspace flow of N_2_ (100 mL/min) was used during the whole cultivation to strip excess CO_2_ or O_2_ from the culture.

Static dish cultures (at 37°C, air + 5% CO_2_ atmosphere) were used in parallel as reference for all tested conditions, with the same inoculum, growth media and seeding/feeding regime performed as in the bioreactors.

### Cell characterization

#### Cell count and viability

Cell density was measured in triplicate using an electrical current exclusion method at a size range of 7.5-15 μm and 5-15 μm in expansion and differentiation cultures, respectively (CASY Model TCC; OLS OMNI Life Science; Germany; or Z2 Coulter Counter; Beckman Coulter; Indianapolis, IN). Population fold change (FC) was calculated in reference to cell numbers at the start of the growth experiments: FC = N(t_i+1_)/N(t_0_). Viability was determined using a hemocytometer and a dye exclusion method (Trypan Blue; Sigma).

#### Differentiation and viability measurements using flow cytometry

About 200.000 cells were stained in HEPES buffer + 0.5% BSA (30 minutes, 4°C), measured using an BD FACSCanto™ II or Accuri C6 flow cytometer (BD Biosciences), gated against specific isotypes, and analyzed using FlowJo™ (version 10.3; Ashland, OR). Antibodies or reagents used were: (i) CD235a-PE (1:2500 dilution; OriGene cat#DM066R), CD49d-BV421 (1:100 dilution; BD-Biosciences cat#565277), DRAQ7 (live/dead stain; 1:200 dilution; ThermoFischer Scientific cat#D15106); (ii) CD235a-PE (1:2500 dilution; OriGene cat#DM066R), CD71-APC (1:200 dilution; Miltenyi cat#130-099-219); (iv) PI (live/dead stain; 1:2000 dilution; Invitrogen cat#P3566); (v) AnnexinV-FITC (1:1000 dilution; BioLegend cat#640906), DRAQ7 (live/dead stain; 1:200 dilution; ThermoFischer Scientific cat#D15106); (vi) DRAQ5 (nuclear stain; 1:2500 dilution; abcam cat# ab108410). Stainings with panels (iv), (v) and (vi) were performed 10 minutes before analysis without intermediate washing steps.

#### Cell morphology and hemoglobin content

Cell pellets (2.5×10^6^ cells, after centrifugation for 5 minutes at 600g) were washed with PBS, resuspended in PBS + 5% HSA, and smeared onto microscope slides. Slides were left to air overnight, stained with Hemacolor® Rapid Staining (Sigma), mounted with Entellan® (Merck; Germany), and examined by bright field microscopy (Leica DM5500B, Leica Microsystems, Germany). Hemoglobin content was determined using *o*-phenylenediamine as previously described (Bakker et al., 2004).

#### Lactate and ammonium measurements

Cell culture samples were taken daily before and after medium refreshment. After centrifugation (600g, 5 min), the supernatant was snap frozen in liquid nitrogen and stored at −80 C until further analysis. Upon thawing, lactate concentrations were measured using a RAPIDlab 1265 blood analyzer (enzymatic amperometric biosensor; Siemens Healthineers, Germany). Ammonium concentration was determined via an enzymatic spectrophotometric assay (Sigma cat#AA0100). Average daily growth rate (μ_max_), doubling time (t_1/2_), and the cell-specific metabolite consumption or production rates (q_lac_, q_NH4_) were calculated as described in Supplemental Methods.

#### Oxygen consumption rate determination

For the determination of the oxygen consumption rate of proliferating erythroblast, bioreactor cultures were run at 200 rpm, with no sparging (no active pH or dO_2_ control), and with a fixed overhead flow of 100 mL/min air + 5% CO_2_ as only source of oxygen. pH was continuously monitored, and passively controlled by the equilibrium between CO_2_ in the overhead and the sodium bicarbonate present in the culture medium (pH variation between 7.2 and 7.7 during the cultivation). The cell-specific consumption rate of oxygen (q_O2_) for each day of culture was determined by the dynamic method, fitting the measured dO_2_ data as described in Supplemental Methods and using the experimentally determined mass transfer coefficient of the system (k_L_a = 0.82 1/h) at the culture conditions.

### Statistical analysis

Statistical analyses were performed using unpaired two-tailed two-sample equal-variance Student’s *t*-test. All data in figures represent the mean ± the standard deviation of the measurements. The number of replicates is N=3 for all experiments, unless indicated.

## Results

### Oxygen consumption by proliferating erythroblasts

To determine the feasibility of erythroblast proliferation in stirred tank bioreactors (STRs), day 9 PBMC-derived erythroblast cultures were seeded in a 2 L STR (working volume = 1 L). Cells were cultured with a stirring speed of 50 rpm (marine impeller; tip speed = 118 mm/s), with air diffusion from the headspace as the sole source of oxygen for the cell culture, similarly to aeration in static culture dishes and orbital shaking (Aglialoro et al., 2021; Heshusius et al., 2019). We observed complete depletion of oxygen after 6-12 hours in the bioreactor culture, followed by a decrease in cell growth and viability compared to a parallel culture in dishes, albeit only after 24 hours (Figures 1A-C; representative experiment). Continuous sparging of air + 5% CO_2_ caused excessive foam production, with cell debris accumulating in foam (data not shown). Proliferation and viability were restored to levels similar or better compared to those of static cultures when headspace aeration was complemented with sparging triggered when the measured dissolved oxygen (dO_2_) concentration was lower than 2.88 mg O_2_/L (equivalent to 40% of saturation with air; a setpoint typically used in STR mammalian cell cultures; (Jan et al., 1997; Ozturk & Palsson, 1991; Restelli et al., 2006)), (Figures 1B-C).

**Figure 1.**
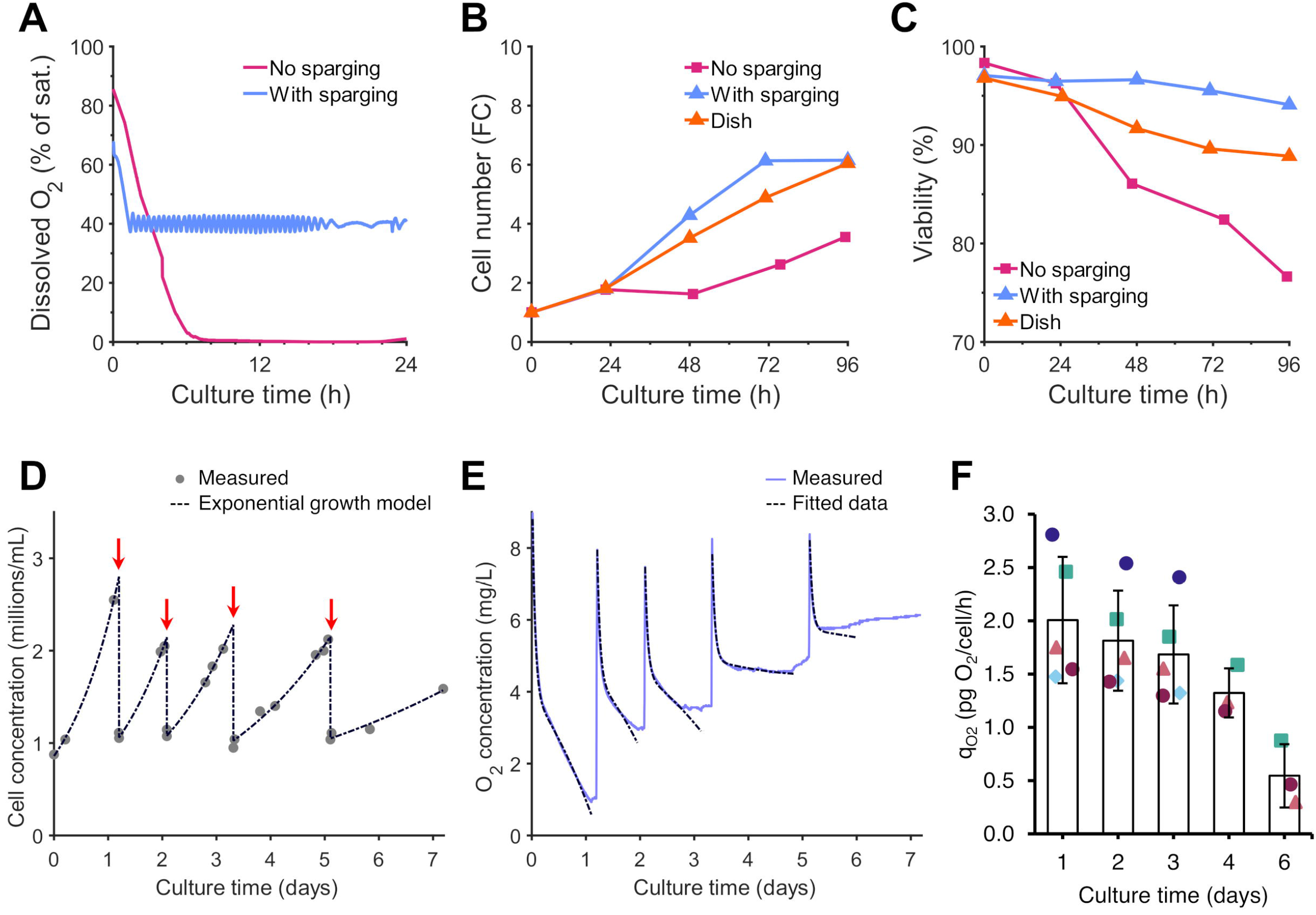
Oxygen can be limiting for erythroblast proliferation in stirred tank bioreactors when headspace aeration is used as sole O_2_ source. **A-C:** Erythroblasts were expanded from PBMCs for 9 days, and subsequently seeded in culture dishes (orange line) or in a 1.5 L STR (working volume: 1 L; stirring speed: 50 rpm; marine down-pumping impeller, diameter: 4.5 cm; 37°C), in which oxygen was provided by gas flow (air + 5% CO_2_) in the headspace (0.3 L/min; purple line, no sparging), or by intermittent sparging triggered at <40% dissolved oxygen (dO_2_: % of the oxygen saturation concentration at the culture conditions; blue line), while pH (∼7.20) was maintained by bicarbonate-buffered medium in equilibrium with the headspace gas. (A) Dissolved oxygen concentration in the culture was measured continuously. (B) Erythroblast cell concentration was monitored daily, with medium refreshment if the measured cell concentration was >1.2×10^6^ cells/mL. Fold change (FC) in cell number was calculated relative to the number of erythroblasts at the start of culture. (C) Culture viability was determined using a trypan blue dye exclusion method. Panels A-E show data for a representative reactor run. D-F: To determine the oxygen requirements of proliferating erythroblasts, day 9 cells were inoculated in 0.5 L STRs (working volume: 300 mL; stirring speed: 200 rpm; marine down-pumping impeller, diameter: 2.8 cm), with a headspace flow of 100 mL/min (air + 5% CO_2_) as only source of oxygen for the culture. (D) A constant growth rate was fitted for each time interval between consecutive medium refreshment events. (E) The drop of dissolved oxygen after each medium refreshment was used to estimate the cell-specific oxygen consumption rate (q_O2_) for each time interval, using the experimentally determined mass transfer coefficient (k_L_a) of 0.82 1/h (see Supplemental Methods). (F) The average cell-specific oxygen consumption rate, q_O2_, during erythroblast expansion was calculated as the mean of at 5 independent bioreactor runs (time intervals for which the q_O2_ was calculated for each run available in Supplemental Figure S1).

The results prompted to measure the oxygen consumption, for which smaller culture volumes were used (in standardized 0.5 L bioreactors) with headspace aeration (100 mL/min of air + 5% CO_2_) and an increased stirring speed (200 rpm; marine impeller; tip speed = 293 mm/s). Day 9 expansion cultures, established from PBMCs, were measured throughout 6 subsequent days of proliferation. Cell numbers were assessed daily, and cultures were diluted with fresh medium, transiently increasing the oxygen concentration (Figures 1D-E). A wide variation of dO_2_ was observed, going from as high as 100% of saturation after medium refreshment, to as low as 10% when high cell concentrations were reached. Oxygen was not limiting in these cultures, due to the higher mass transfer coefficient (k_L_a = 0.82 1/h), and the lower total oxygen demand because of the lower culture volume. The cell-specific oxygen consumption rate (q_O2_) was calculated for 5 independent cultures at distinct days (Supplemental Methods) and decreased from 2.01±0.53 at the start of the experiment (day 9 of culture) to 0.55±0.24 pg/cell/h 6 days later (Figure 1F; Supplemental Figure S1). The decrease in oxygen consumption mirrored a decrease in cell proliferation (Figure 1D), possibly due to enhanced spontaneous differentiation of erythroblasts.

So, although headspace aeration can ensure oxygen availability in 0.5 L bioreactor cultures, a better aeration strategy is required to ensure sufficient O_2_ supply in cultures with increased volumes, and to evaluate the effect of defined constant dO_2_ levels in cultured erythroblasts.

### Controlled aeration supports erythroblast expansion in stirred tank bioreactors

To improve the dO_2_ control in 0.5 L bioreactors, we used intermittent sparging of air, combined with a continuous headspace flow of nitrogen (100 mL N_2_/min) to strip excess O_2_ from the culture and drive dO_2_ to the desired setpoint (40% dO_2_) (Supplemental Figure S2A). Intermittent sparging of CO_2_ controlled the pH (=7.5; Supplemental Figure S2B).

Day 9 PBMC-derived erythroblast cultures were seeded in the bioreactors at an initial cell concentration of 0.7×10^6^ cells/mL (Figure 2A). Erythroblasts seeded in culture dishes were cultured in parallel using similar medium refreshment conditions as reference.

**Figure 2.**
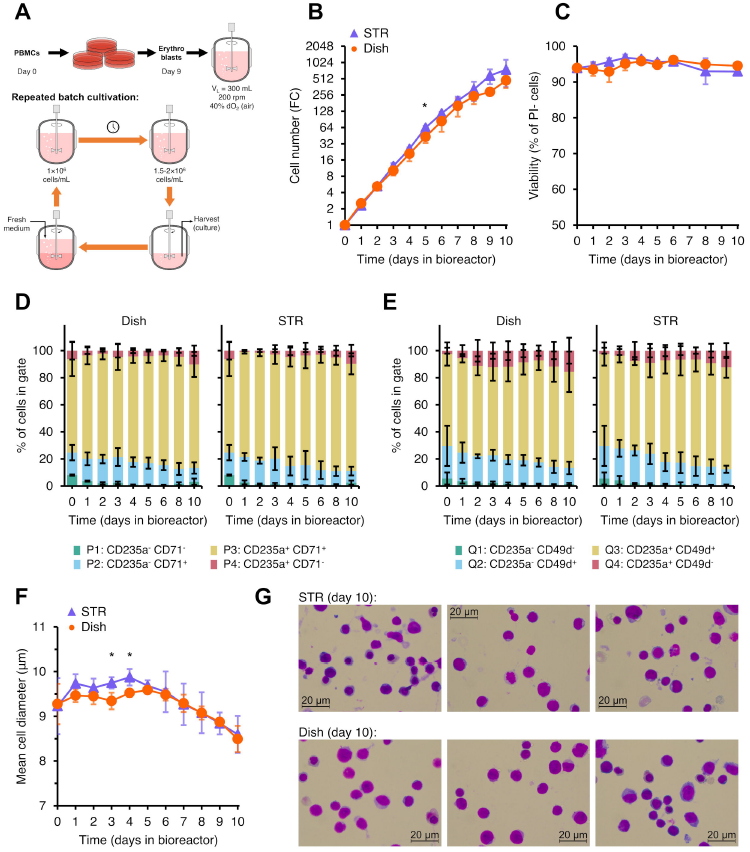
Efficient expansion of erythroblasts can be achieved in stirred tank bioreactors. Erythroblasts were expanded from PBMCs for 9 days, and subsequently seeded in culture dishes (orange lines) or STRs (blue lines) at a starting cell concentration of 0.7×10^6^ cells/mL. STRs were run with a constant N_2_ headspace flow to fully control dO_2_ and pH. **(A)** Cells were cultured using a sequential batch feeding strategy: medium was refreshed when the cell concentration (measured daily) was >1.2×10^6^ cells/mL. **(B)** Cell concentration during 10 days of expansion (fold change (FC) compared to start of the culture). **(C-E)** Cells were stained with propidium iodide (PI; the percentage of PI^-^ cells indicate the % of viable cells) **(C)**, and with CD235a plus CD71 **(D)**, or CD235a plus CD49d **(E)** (gating strategy available in Supplemental Figure S3). **(F)** Mean cell diameter (of cells >5 µm) was measured daily. **(G)** Cytospin cell morphology by May-Grünwald-Giemsa (Pappenheim) staining of cultures after 10 days of expansion. All data is displayed as mean ± SD (error bars; n=3 reactor runs / donors). Significance is shown for the comparison with dish cultures (unpaired two-tailed two-sample equal-variance Student’s *t*-test; \**P*<0.05, \*\**P*<0.01, \*\*\**P*<0.001, not displayed if difference is not significant).

The growth profile was comparable between bioreactor and static cultures, showing a 750-fold increase in cell number after 10 days of culture (resp. day 9 to day 18 counting from PBMC isolation and seeding; Figure 2B). No significant difference was found between the viability of bioreactor and dish cultures (day 10 PI^-^ events: 92.9%±1.1% in bioreactors, 94.5%±0.4% in culture dishes; Figure 2C). Flow cytometry indicated that most cells were committed to the erythroid lineage at the start of the bioreactor culture (<10% CD235a^-^/CD71^-^ cells; >70% CD235a^+^; Figure 2D; gating strategy available in Supplemental Figure S3). Although sustained proliferation of CD235a^+^/CD49d^+^/CD71^high^ erythroblasts can be maintained in presence of EPO, hSCF and dexamethasone, some spontaneous differentiation can take place, initially leading to an increase in CD235a^+^ cells, and a subsequent gradual decrease of CD49d and CD71 expression. Both in bioreactors and dishes the percentage of CD235a^+^/CD71^low^ and CD235a^+^/CD49d^-^ cells was maintained lower than 10% during the whole cultivation (Figures 2D-E). During the last 5 days of the experiment (day 14-19 after seeding PBMCs), the mean cell diameter gradually decreased (Figure 2F). Staining with AnnexinV and DRAQ7 indicated only a low percentage of apoptotic or dead cells (Supplemental Figure S3D), whereas cytospins indicated a large portion of cells with condensed nuclei (Figure 2G), indicative of spontaneous differentiation. Thus, spontaneous differentiation, rather than decreased viability, reduced the proliferative capacity of day 17-19 cultures. Importantly, this was similar between static cultures and cells cultured in bioreactors.

### Effect of dissolved oxygen concentration on erythroblast expansion cultures

Standard mammalian cell culture conditions are mostly hyperoxic compared to their native *in vivo* niche, potentially leading to oxidative stress and impaired growth (Mas-Bargues et al., 2019). The 40% dO_2_ used in our initial experiments, equivalent to 2.88 mg O_2_/mL, is 5-fold higher than the oxygen concentration in the bone marrow compartment in which erythroblast proliferation and differentiation takes place (0.6 mg O_2_/mL; (Spencer et al., 2014)). Therefore, we tested whether BM mimicking oxygen concentrations also supported erythroblast expansion. Bioreactor cultures at 10% dO_2_ (0.72 mg O_2_/L) showed comparable cell yields to dish conditions (Figure 3A), while requiring lower volumes of sparged air compared to 40% dO_2_ (Supplemental Figure S4A). A continuous decrease of growth rate, from 0.78±0.19 1/day to 0.45±0.05 1/day after 8 days of culture, was observed in bioreactor cultures at 10% dO_2_. By contrast, in hyperoxic bioreactor cultures (dO_2_ = 40%) the growth rate was stable at ∼0.78 1/day for the first 5 days of culture, followed by a decrease to 0.38±0.12 1/day at day 8 of cultivation (Figure 3B; growth rates calculated using cell counts available in Supplemental Figure S4C).

**Figure 3.**
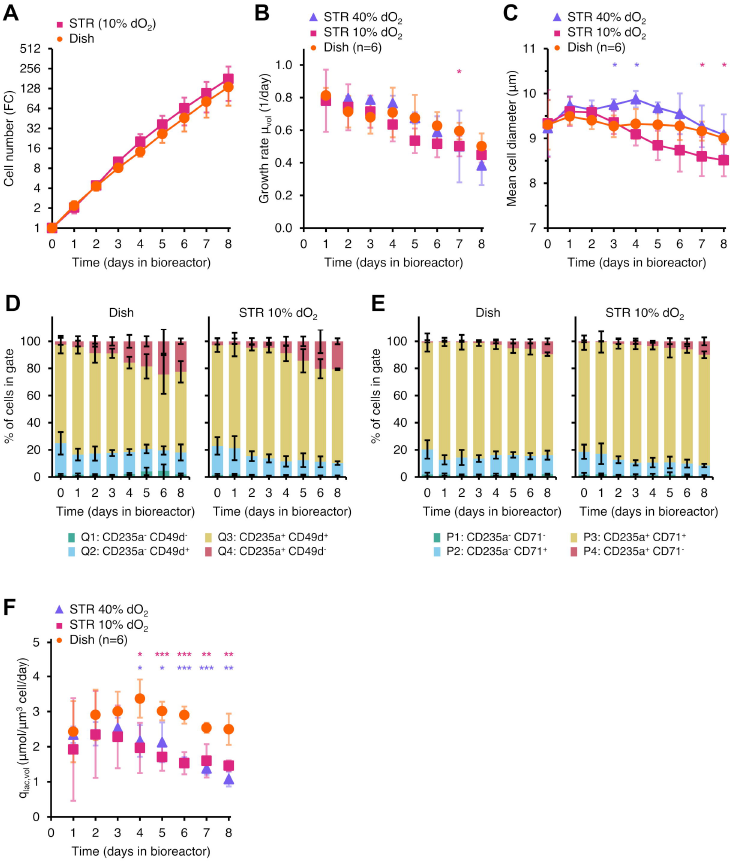
Low dissolved oxygen concentration support erythroblast expansion. Erythroblasts were expanded from PBMCs for 9 days, and subsequently seeded in culture dishes (orange line) or STRs (100 mL/min N_2_ headspace flow) at a starting cell concentration of 0.7×10^6^ cells/mL. Dissolved oxygen was controlled by air sparging when below the targeted setpoint (10% (purple line) or 40% (blue line), equivalent to 0.72 and 2.88 mg O_2_/L respectively), and by stripping of excess oxygen using 100 mL/min N_2_ headspace flow. **(A)** Cell concentration monitored for 8 days of culture, with medium refreshment when the measured cell concentration (daily) was >1.2×10^6^ cells/mL. Fold change (FC) in cell number was calculated using the number of erythroblasts at the start of culture. **(B)** Growth rate was calculated for each day using the total biomass concentration (μm^3^ of total cell volume per mL of culture) assuming exponential growth between consecutive media refreshment events. **(C)** Mean cell diameter was measured daily. (**D-E)** Cells were stained with CD235a plus CD71 **(D)**, or CD235a plus CD49d **(E)**. **(F)** Cell-specific lactate production rate (q_lac,vol_) calculated using growth rate data and measured extracellular lactate concentrations (see Supplemental Methods). All data is displayed as mean ± SD (error bars; n=3 reactor runs / donors, unless indicated otherwise). Significance is shown for the comparison with dish cultures (unpaired two-tailed two-sample equal-variance Student’s *t*-test; \**P*<0.05, \*\**P*<0.01, \*\*\**P*<0.001, not displayed if difference is not significant). Growth rates and q_lac_ calculated using cell counts available in Supplemental Figure S4.

Cell size also decreased faster during the culture period at 10% dO_2_ (Figure 3C). This may indicate partial differentiation within the population of CD235a^+^/CD71^+^/CD49d^+^ population, or a reduced protein synthesis at this lower O_2_ concentration (Brugarolas et al., 2004). Although the percentage of CD235a^+^/CD49d^-^/CD71^low^ cells increased during culture, this was similar between bioreactors at 10% dO_2_ and standard static cultures (Figure 3D-E). Lactate, typically produced by aerobic glycolysis and glutaminolysis, may negatively influence erythroblast growth and viability. Extracellular lactate concentrations were typically <6 mM during culture in bioreactors or culture dishes (Supplementary Figure S2E). Nevertheless, a consistently lower cell-specific lactate production rate (q_Lac,vol_) was observed in bioreactor conditions both at 10 and 40% dO_2_, compared to culture dishes (>20% reduction; Figure 3F; q_Lac,counts_ available in Supplemental Figure S4D), suggesting a higher rate of aerobic glycolysis in static cultures. The q_Lac_ values decreased after 4 days of culture in both bioreactors and dish cultures to the same extent, which may be due to the concurrent reduction in growth rate. In addition to lactate, ammonia is a common inhibitor of cell proliferation. The ammonia production, however, was similar in bioreactor runs at 10% or 40% dO_2_ and in standard dish cultures (Supplemental Figure S4B,E).

Although a 10% dO_2_ setpoint is closer to the physiological oxygen concentrations in which erythroblasts proliferate *in vivo*, we conclude that the lower dO_2_ does not alter growth or spontaneous differentiation compared to a 40% dO_2_ setpoint.

### Shear stress affects erythroblast proliferation during expansion cultures

Shear stress is another critical parameter in mammalian cell cultures. It can have a negative effect on growth and viability (Neunstoecklin et al., 2015), whereas insufficient stirring speeds leads to inadequate mixing and aeration when scaling up (Xing et al., 2009). To evaluate the effect of stirring speed on erythroblast proliferation, erythroblasts were inoculated in 0.5 L bioreactors, and the impeller speed was increased from 200 rpm to 600 rpm (tip speed = 880 mm/s) and 1800 rpm (tip speed = 2640 mm/s). After 6 days of culture, a 87.2±2.7-fold change in cell number was observed in the 600 rpm cultivations (Figure 4A), similar to the growth levels previously observed at 200 rpm and in static cultures. By contrast, an agitation speed of 1800 rpm reduced proliferation during the first 4 days of culture, which, however, recovered to values comparable to those observed at 600 rpm after 6 days of culture (Figure 4B).

**Figure 4.**
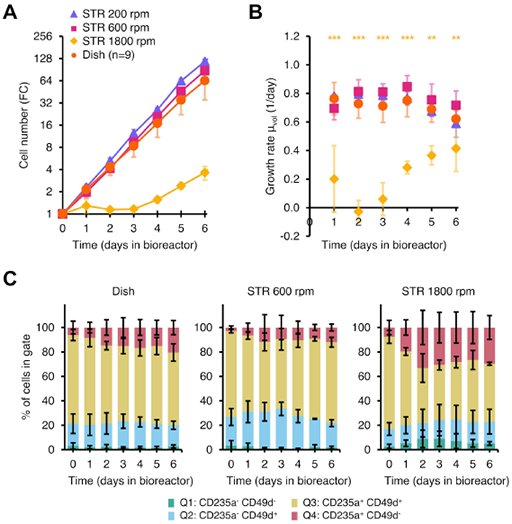
High stirring speeds can sustain erythroblast expansion. Erythroblasts were expanded from PBMCs for 9 days, and subsequently seeded in culture dishes (orange line) or STRs (dO_2_: 40% controlled by sparging of air; 100 mL/min N_2_ headspace flow) at a starting cell concentration of 0.7×10^6^ cells/mL, under agitation at 200 (blue line), 600 (purple line) or 1800 rpm (yellow line). **(A)** Cells were maintained between 0.7 and 1.5×10^6^ cells/mL by dilution with fresh medium. Cumulative cell numbers were calculated and represented as fold change (FC) compared to the start of the experiment. **(B)** Growth rate for each day was calculated assuming exponential growth between consecutive media refreshment events. **(C)** Cells were stained with CD235a plus CD49d to evaluate the progression of spontaneous differentiation during culture. All data is displayed as mean ± SD (error bars; n=3 reactor runs / donors, unless indicated otherwise). Significance is shown for the comparison with dish cultures (unpaired two-tailed two-sample equal-variance Student’s *t*-test; \**P*<0.05, \*\**P*<0.01, \*\*\**P*<0.001, not displayed if difference is not significant).

The increase in stirring speed to 600 rpm did not enhance spontaneous differentiation, with only 12.1±2.8% cells in the most differentiated CD235^+^ CD49d^-^ compartment at day 6. At 1800 rpm, however, CD49d expression decreased in the first 3 days of culture, stabilizing in the following days (Figure 4C). Cell cycle progression was not affected by the stirring speed (Supplemental Figure S5).

### Terminal differentiation of erythroblast cultures in bioreactors

We validated that erythroblasts expanded in bioreactors could be differentiated as standard static cultures throughout all conditions (data not shown). To examine terminal differentiation in bioreactors, day 10 PBMC-derived erythroblast cultures were seeded in differentiation medium at a starting cell concentration of 1.5×10^6^ cells/mL, both in bioreactors and culture dishes (Figure 5A). Bioreactors were controlled with the same conditions as in the reference expansion phase (40% dO_2_, pH 7.5, 200 rpm, 37°C). During the first 3 days of differentiation, cell numbers increased in both systems. Simultaneously, sparging was required to maintain dO_2_ at 40% (Supplemental Figure S6). This was followed by a cell cycle arrest in both static and bioreactor conditions. Bioreactor cultures showed a gradual decrease in cell numbers until the end of culture, while cell numbers in static conditions remained similar (Figure 5B). Concurrently, the dO_2_ concentration increased in absence of sparging after 4 days of culture.

**Figure 5.**
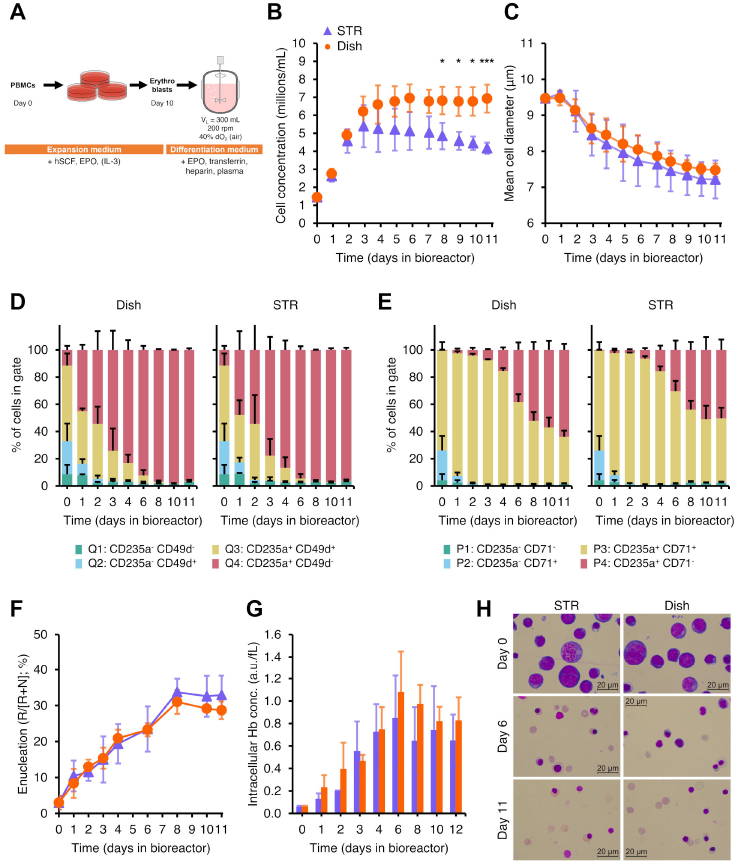
Erythroblast differentiation can be achieved in stirred tank bioreactors. Erythroblasts were expanded from PBMCs for 10 days, and subsequently seeded in differentiation medium at a starting cell concentration of 1×10^6^ cells/mL. **(A)** Cells were transferred to culture dishes or STRs and kept in culture for 11 subsequent days without medium refreshment. **(B)** Cell concentration during 11 days of differentiation in STRs (blue symbols) and dishes (orange symbols). **(C)** Mean cell diameter was measured daily. **(D-E)** Cells were stained with CD235a plus CD49d **(D)**, or CD235a plus CD71 **(E)**, and percentages in each quadrant are shown. **(F)** Enucleation percentage of erythroid cells was calculated from the forward scatter and DRAQ5 staining. DRAQ5^-^ cell numbers (reticulocytes, R) were divided by the sum of small DRAQ5^+^ events (nuclei, N + reticulocytes, R). **(G)** Hemoglobin was measured in arbitrary units (a.u.) and the intracellular hemoglobin concentration was calculated using the total cell volume. **(H)** Representative cytospin cell morphology by May-Grünwald-Giemsa (Pappenheim) staining of bioreactor cultures during differentiation. All data is displayed as mean ± SD (error bars; n=3 reactor runs / donors). Significance is shown for the comparison with dish cultures (unpaired two-tailed two-sample equal-variance Student’s *t*-test; \**P*<0.05, \*\**P*<0.01, \*\*\**P*<0.001, not displayed if difference is not significant).

Differentiation was evident from a decrease in mean cell diameter in bioreactors and culture dishes (Figure 5C) with a concomitant loss of CD49d and CD71 expression (Figure 5D-E). Hemoglobin accumulation and enucleation efficiency was also similar in static and stirred bioreactors (Figure 5F-G). Presence of enucleated cells and egressed nuclei was confirmed by cytospins (Figure 5H). In conclusion, erythroblast differentiation could also be supported in our stirred bioreactor systems, reaching similar levels of enucleation to those of culture dishes, albeit with slightly lower cell yields.

### Expansion cultures can be scaled up to 3 L bioreactors

Knowing the boundaries to cultured erythroblast in STRs enabled us to scale-up from 300 mL cultures in 500 mL STRs to larger volumes. Day 8 erythroblasts cultures from PBMCs were seeded at a density of 0.7×10^6^ cells/mL in a 500 mL STR (initial culture volume = 100-150 mL), and cell density was adjusted daily to 0.7×10^6^ cells/mL by adding more medium. When the volume surpassed 400 mL, cells were transferred to a 3 L STR (minimum working volume = 800 mL), operated with a tip speed similar to that tested in 0.5 L bioreactors (300 mm/s; single down-pumping marine impeller; diameter = 5 cm; 115 rpm). Medium was added to maintain cell density until a total volume of 2.5 L was reached. Subsequently, the total volume was maintained at 2.5 L and excess cells were removed (Figure 6A-B). The cell cultures proliferated exponentially for 9 subsequent days achieving an average 196-fold increase in the 3L STR, and 234-fold under static conditions (Figure 6C). Interestingly, the variation between 3 cultures was smaller in the STR (range: 112-280-fold) compared to the same cultures expanded in dishes (range: 51-462-fold). After 9 days of culture, there was no difference in viability or differentiation stage of the cells (Figure 6D-E; Supplemental Figure S7A). Similar to the 300 mL cultures, lactate production in 2.5 L STRs was lower compared to static conditions (Supplemental Figure S7B).

**Figure 6.**
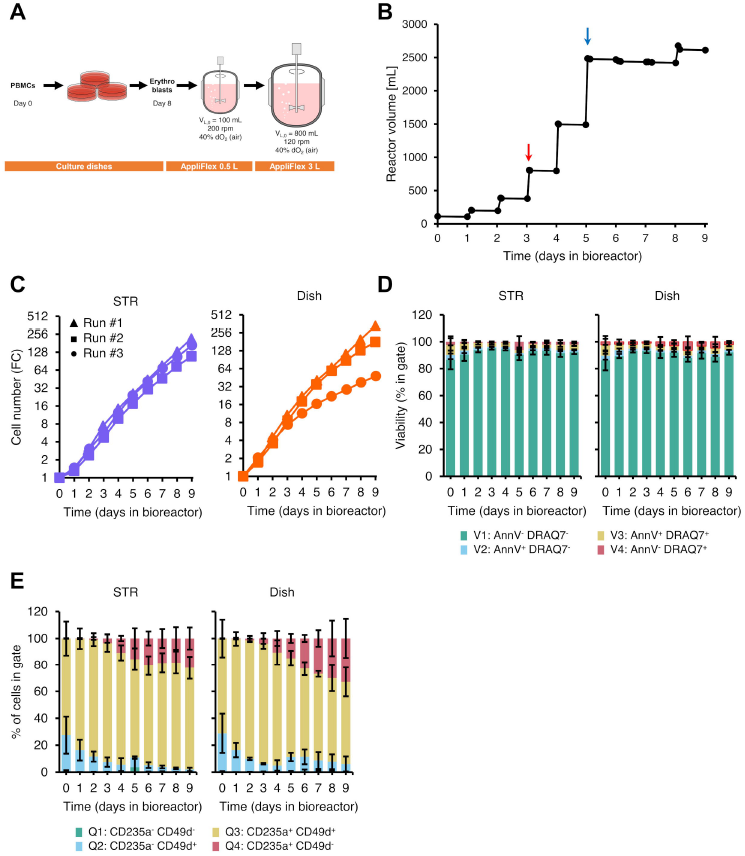
Scale-up of erythroblast expansion to 3 L stirred tank bioreactors. Erythroblasts were expanded from PBMCs for 8 days, and subsequently seeded in culture dishes or 0.5 L stirred tank bioreactors (starting volume: 100-200 mL) at a starting cell concentration of 0.7×10^6^ cells/mL. **(A)** Cells were kept in culture following a fed batch feeding strategy in which medium was refreshed if the measured cell concentration (daily) was >1.2×10^6^ cells/mL. Upon reaching a total number of >400 million cells, the culture was transferred to a 3.0 L bioreactor (starting volume: 800 mL; 115 rpm with marine down-pumping impeller, diameter: 5.0 cm), which was progressively filled by daily medium additions. **(B)** Culture volume in the bioreactor for an exemplary run. Transition from the 0.5 L to the 3 L reactor was performed at day 3 of culture (red arrow). Upon filling of the 3 L reactor (blue arrow), excess cells were harvested daily to keep a working volume of 2.5-2.7 L. **(C)** Erythroblast cell concentration was monitored for 9 days of culture. Fold change (FC) in cell number was calculated relative to erythroblast numbers at the start of culture. The same preculture was seeded in parallel in the STR and in a dish. **(D-E)** Cells were stained with AnnexinV (apoptosis staining) and DRAQ7 (cell impermeable DNA stain) **(D)**, or CD235a plus CD49d (erythroid differentiation markers). Percentage of cells in each quadrant is included. **(E)**. All data is displayed as mean ± SD (error bars; n=3 reactor runs / donors).

## Discussion

Efficient cRBC production requires the transition from static culture systems to scalable cultivation platforms. We established the process conditions (oxygen and agitation) required for effective expansion and differentiation of erythroblast cultures in STRs. Oxygen availability was critical during expansion of erythroblast cultures, whereas the requirement for oxygen decreased during terminal differentiation. Stirring speeds required for a homogeneous distribution of cells in the bioreactor did not affect cell growth or differentiation. Only much elevated stirring speeds led to a temporary cell growth arrest and an acceleration in spontaneous erythroblast differentiation. Within the operating boundaries established in this study, the STR culture volume could be scaled up from 300 mL to 2500 mL.

### Oxygen requirements of erythroblast in culture

Using a combination of headspace N_2_ gas flow and intermittent air allowed us to tightly control dO_2_ in our bioreactor cultures. While low dO_2_ setpoints can reduce the overall sparging requirements of the culture and are closer to the *in vivo* hematopoietic niche, no significant improvement on growth, lactate accumulation or spontaneous differentiation was observed in our experiments when the dO_2_ setpoint was decreased from 40% to 10%. Although oxygen concentration setpoints lower than 10% dO_2_ may enhance proliferation, maintaining a constant dO_2_ <10% in our set-up is technically challenging due to the noise in dO_2_ readings and the oscillations caused by the discrete sparging events. This could be addressed by using oxygen probes with lower response times or with aeration strategies in which gas composition is adjusted based on the dO_2_ measurements, avoiding large oxygen concentration peaks during sparging. Studying the effect of lower and fluctuating dO_2_ concentrations can provide further information on the potential challenges that could be faced in the large bioreactors required for the production of the required number of cRBCs for a single transfusion unit, in which oxygen and nutrient gradients are difficult to avoid, and the risk of having oxygen-limited regions is higher (Anane et al., 2021; Serrato et al., 2004).

The oxygen requirements of erythroblasts decreased during culture, from 2.01 to 0.55 pg/cell/h, and increasingly less during terminal differentiation. These values explain the fast decrease in dO_2_ concentration in the initial 2 L bioreactor experiments, in which the k_L_a (0.27 1/h; headspace aeration only) could only support theoretical early erythroblast concentrations of <0.97×10^6^ cells/mL. Seahorse assays, typically used to quantify the oxygen consumption rates of erythroblasts, are performed with low cell numbers and in an assay medium with a composition different to that used for culture, potentially resulting in q_O2_ values not representative of the culture conditions (van der Windt et al., 2016). Bayle et al. reported much lower q_O2_ values for their bioreactor cultures, decreasing from 0.063 pg/cell/h for early erythroblasts to 0.017 pg/cell/h in the later stages of differentiation (Bayley et al., 2017). A similar decrease in oxygen requirements during erythroid differentiation was described by Browne et al., from 2.84 to 0.13 pg/cell/h for proerythroblasts and reticulocytes, respectively (Browne et al., 2014). This decrease in oxygen requirement could be a result of mitophagy at terminal stages of erythroblast differentiation decreasing mitochondria numbers and marking the shift to anaerobic metabolism as occurring in RBCs (Barde et al., 2013; Chen et al., 2008; Sandoval et al., 2008). Differences between our measured q_O2_ values and those previously reported could be explained by differences in the differentiation status of erythroid cells used at the start of the bioreactor cultures, culture medium composition or oxygen concentration (Moradi et al., 2021). Of note, our values are in agreement with measured oxygen uptake rates of other human cell lines (0.4-6.2 pg O_2_/cell/h; (Wagner et al., 2011)). We suggest using continuous dO_2_ measurements and the dynamic method to estimate q_O2_ in erythroid cultures, as other methods such as using offgas data are difficult to implement due to the low oxygen consumption rates of mammalian cell cultures (Singh, 1996).

### Erythroid expansion cultures are robust to high stirring speeds

This study shows that erythroblast expansion cultures can withstand agitation speeds of up to 600 rpm (tip speed = 880 mm/s), without negatively affecting growth, viability or the differentiation stage of the cells. Surprisingly, very high stirring speeds (1800 rpm) led to a temporary arrest in growth, followed by a recovery after 5-6 days, accompanied by an acceleration in erythroid differentiation as reported previously for erythroid cultures in flasks under orbital shaking (Aglialoro et al., 2021). This recovery in growth could be explained by an adaptation of culture cells to these turbulent conditions, or by the selection of erythroblast cells more tolerant to agitation during the first days of culture. A more thorough characterization of this adaptation on not only the erythroblast transcriptome but also at the lipidomic and metabolomic level will need to be performed to identify (signaling) pathways triggered by this mechanical stress, the nature of the adaptative response (e.g. membrane composition remodeling), and to determine the mechanism under which erythroid differentiation is promoted under these conditions.

Interestingly, other studies have reported less tolerance of erythroblast cultures to agitation. Increased apoptosis, acceleration of erythroid differentiation, and lower enucleation levels have been observed in a gyro-rocker bioreactor at low orbital speeds (20 rpm) (Boehm et al., 2010). Bayley et al. also observed a decrease in proliferation in stirred bioreactors compared to static cultures, although no significance difference on growth was observed between stirring speeds of 300 and 450 rpm (tip speeds = 180 and 270 mm/s, respectively) (Bayley et al., 2017). Han et al. observed a dependence on the inoculum age to agitation tolerance, with proerythroblast and basophilic erythroblast cultures showing a quick decrease in growth and viability in microbioreactors agitated at 300 rpm (tip speed = 180 mm/s), while more mature cells (day 12 after CD34^+^ isolation and start of culture), could tolerate the same conditions (Han et al., 2021). However, it is difficult to directly compare the hydrodynamic conditions between these reports, partly due to differences in the culture set-up but also due to a lack of consensus on which mixing parameter (rpm, tip speed, volumetric power input) is more appropriate to do this comparison, which together co-define the dynamic shear stress experienced by cells. Local turbulent energy dissipation rate (EDR) has been suggested as an alternative parameter to define optimal ranges of agitation for mammalian cell culture (Chalmers, 2015). Supporting the relevance of energy dissipation rate, it has been recently reported that turbulence and not shear stress is critical for large scale platelet production (Ito et al., 2018). Estimates of EDR in STRs could be useful to rationally drive the scale-up of the process, ensuring limited exposure of erythroblast to excessive hydrodynamic forces.

Although the effect of shear stress and EDR can be evaluated in STRs by varying the agitation rate, changes in stirring speed also affect the mass transfer rate between the gas and liquid phases, impacting the aeration requirements to control dO_2_ and pH at the defined setpoints. This, in turn, can influence the exposure of cells to high local O_2_ (and CO_2_, if pH is controlled with this gas) concentrations and to high EDR regions due to bubble bursting phenomena (Walls et al., 2017). A full uncoupling of agitation and aeration is challenging, as other aeration strategies such as using submerged oxygen-permeable membranes, can also lead to undesired effect such as very high local O_2_ concentrations (Aunins & Henzler, 2001; Côté et al., 1989).

### Quantification of metabolite consumption/production rates in erythroid cultures

We observed a decrease in erythroblast proliferation during expansion cultures, with growth rates as low as ∼0.3 1/day at the end of the cultures. Growth limitations in erythroblast batch and fed-batch cultures has been previously reported, but the origin of this limitation has not been identified yet (Bayley et al., 2017; Glen et al., 2018). Here we report lactate and ammonium concentrations below 6 and 0.6 mM respectively, which are lower than typical growth-inhibiting concentrations for other animal cell lines (Cruz et al., 2000; Hassell et al., 1991; Ozturk et al., 1992). Interestingly, we measured lactate production rates (q_Lac_) of 0.5-2.5 pmol/cell/day, significantly lower than those reported for other erythroid cultures (3-12 pmol/cell/day; (Bayley et al., 2017; Lee et al., 2018; Patel et al., 2000; Sivalingam et al., 2020)). Dissolved oxygen concentration in the bioreactor did not have a significant effect on q_Lac_, but static cultures consistently showed higher rates in later days of culture. Oxygen limitations caused by low oxygen concentrations (<10% dO_2_) in the bottom of culture dishes could lead to an increase in anaerobic glycolysis rates in erythroblasts cultured under static conditions, leading to the observed lactate production (Al-Ani et al., 2018). Even at the measured low concentrations, lactate can potentially induce activation of erythroid-related genes such as STAT5, affecting erythroid differentiation dynamics (Luo et al., 2017). Strategies to limit lactate production in culture could be implemented, such as feeding profiles that limit glucose concentrations during culture.

### Outlook

A 200-fold expansion per PBMC already takes place in the first 8 days of culture (before reactor inoculation), still performed under static conditions in culture dishes (Heshusius et al., 2019), resulting in an overall 200×750 = 150.000- fold expansion after 10 days of culture in the bioreactor. The high cell yields in this first culture stage are partly due to the positive contribution of CD14^+^ monocytes/macrophages present in the PBMC pool by the production of soluble factors that support erythroblast expansion (Heideveld et al., 2015; van den Akker et al., 2010). Full translation of our protocol to STRs would require defining operating conditions that would support HSC proliferation and macrophages in the first days of culture.

While we could scale-up erythroblast expansion cultures from dishes to 0.5 L and 3.0 L bioreactors, the large number of cells required for a single cRBC transfusion unit would still require prohibitively large STRs. Perfusion approaches could be used to increase cell concentrations, while continuously removing lactate and other produced inhibitory metabolic byproducts. The cell-retention strategies required for perfusion could also be of value to perform the required medium changes when transitioning from proliferation to differentiation in culture, where removal of hSCF is needed for efficient maturation and enucleation. This, combined with the development of low cost media formulations, would allow for a large scale cost-effective production of cRBCs.

## Abbreviations

BM: Bone marrow
BSA: Bovine serum albumin
CD235a: Glycophorin A
CD49d: Integrin alpha 4
CD71: Transferrin receptor 1
cRBC: Cultured red blood cell
Dex: Dexamethasone
EDR: Energy dissipation rate
Epo: Erythropoietin
EpoR: Erythropoietin receptor
FC: Fold change
HEPES: 4-(2-hydroxyethyl)-1-piperazineethanesulfonic acid
HIF1α: Hypoxia-inducible factor 1, alpha subunit
HSA: Human serum albumin
HSC: Hematopoietic stem cell
hSCF: Human stem cell factor
NFATC2: Nuclear factor of activated T-cells 2
OXPHOS: Oxidative phosphorylation
PBMC: Peripheral blood mononuclear cell
PBS: Phosphate-buffered saline
PI: Propidium iodide
RBC: Red blood cell
rpm: Revolutions per minute
SCF: Stem cell factor
STAT5: Signal transducer and activator of transcription 5A
STR: Stirred tank reactor
TCA: Tricarboxylic acid

## List of symbols

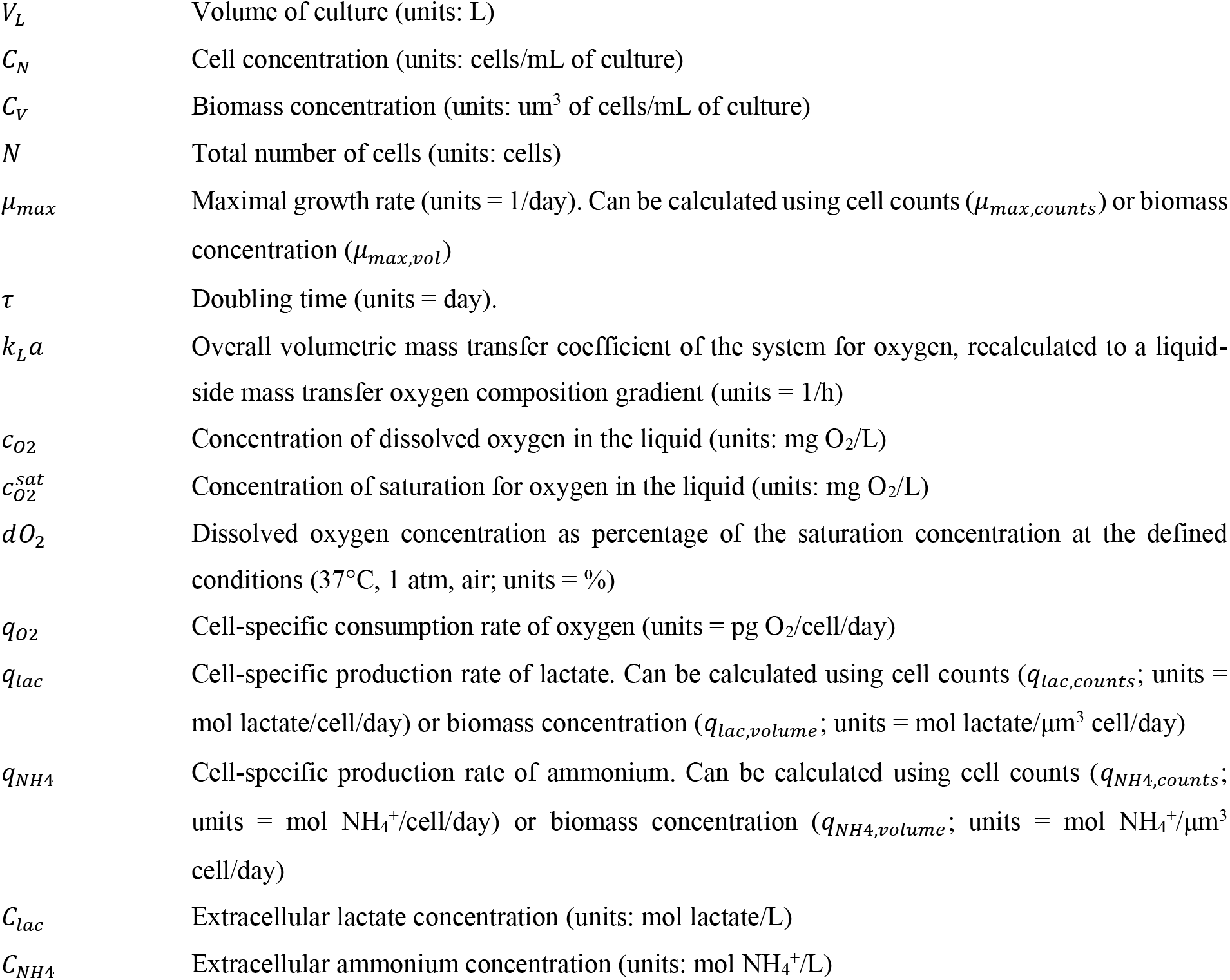

|  |  |
| --- | --- |
| $V_L$ | Volume of culture (units: L) |
| $C_N$ | Cell concentration (units: cells/mL of culture) |
| $C_V$ | Biomass concentration (units: $\mu\text{m}^3$ of cells/mL of culture) |
| $N$ | Total number of cells (units: cells) |
| $\mu_{max}$ | Maximal growth rate (units = 1/day). Can be calculated using cell counts ($\mu_{max,counts}$) or biomass concentration ($\mu_{max,vol}$) |
| $\tau$ | Doubling time (units = day). |
| $k_L a$ | Overall volumetric mass transfer coefficient of the system for oxygen, recalculated to a liquid-side mass transfer oxygen composition gradient (units = 1/h) |
| $c_{O_2}$ | Concentration of dissolved oxygen in the liquid (units: mg $\text{O}_2$/L) |
| $c_{O_2}^{sat}$ | Concentration of saturation for oxygen in the liquid (units: mg $\text{O}_2$/L) |
| $dO_2$ | Dissolved oxygen concentration as percentage of the saturation concentration at the defined conditions (37°C, 1 atm, air; units = %) |
| $q_{O_2}$ | Cell-specific consumption rate of oxygen (units = pg $\text{O}_2$/cell/day) |
| $q_{lac}$ | Cell-specific production rate of lactate. Can be calculated using cell counts ($q_{lac,counts}$; units = mol lactate/cell/day) or biomass concentration ($q_{lac,volume}$; units = mol lactate/$\mu\text{m}^3$ cell/day) |
| $q_{NH_4}$ | Cell-specific production rate of ammonium. Can be calculated using cell counts ($q_{NH_4,counts}$; units = mol $\text{NH}_4^+$/cell/day) or biomass concentration ($q_{NH_4,volume}$; units = mol $\text{NH}_4^+$/$\mu\text{m}^3$ cell/day) |
| $C_{lac}$ | Extracellular lactate concentration (units: mol lactate/L) |
| $C_{NH_4}$ | Extracellular ammonium concentration (units: mol $\text{NH}_4^+$/L) |

## Acknowledgements

We thank Tom van Arragon and Cristina Bernal Martínez (Applikon Biotechnology; Delft; The Netherlands) for technical help and advice on bioreactor cultures. This work was supported by the ZonMW TAS program (project 116003004), by the Landsteiner Foundation for Bloodtransfusion Research (LSBR project 1239), and by Sanquin Blood Supply grants PPOC17-28 and PPOC119-14.

## Authorship

Contribution: J.S.G.M. performed the experiments; J.S.G.M., E.v.d.A., A.W., and M.v.L. designed the experiments, analyzed the data, and wrote the manuscript; G.I. and L.v.d.W contributed to data analysis and writing of the manuscript. All authors critically revised the manuscript.

## Correspondence

Marieke von Lindern, Department of Hematopoiesis, Sanquin Research and Landsteiner Laboratory, Amsterdam UMC, Plesmanlaan 125, 1066CX Amsterdam, The Netherlands; and Joan Sebastián Gallego-Murillo, Department of Hematopoiesis, Sanquin Research and Landsteiner Laboratory, Amsterdam UMC, Plesmanlaan 125, 1066CX Amsterdam, The Netherlands.

