## Supplemental Figures for "Expansion and differentiation of *ex vivo* cultured erythroblasts in scalable stirred bioreactors"

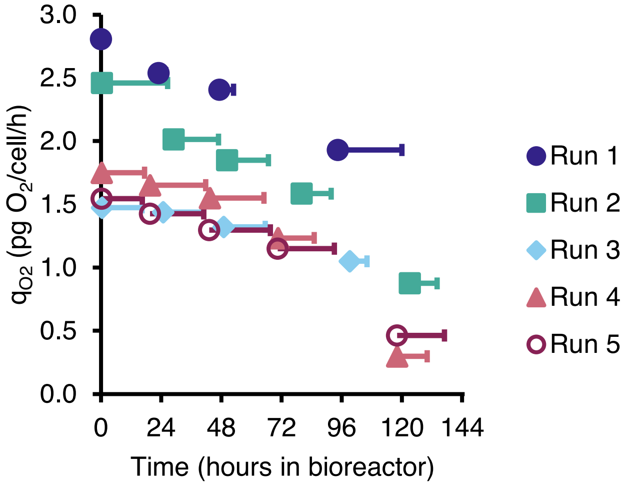


**Supplemental Figure S1. Quantification of oxygen requirements of expanding erythroblasts.** Erythroblasts were expanded from PBMCs for 9 days, and subsequently seeded in 500 mL stirred tank bioreactors (working volume = 300 mL; stirring speed = 200 rpm; marine down-pumping impeller with diameter = 2.8 cm) at a starting cell concentration of 0.7×$10^6$ cells/mL. A headspace flow of 100 mL/min (air + 5% $CO_2$) as only source of oxygen for the culture. Cell-specific oxygen consumption rates ($q_{O2}$) was determined via the dynamic method, using the drop of $dO_2$ after each dilution event and the calculated growth rate between consecutive medium refreshment events (see Supplemental Methods). Length of the bars depict the time interval used for the fitting of the oxygen mass balance model.

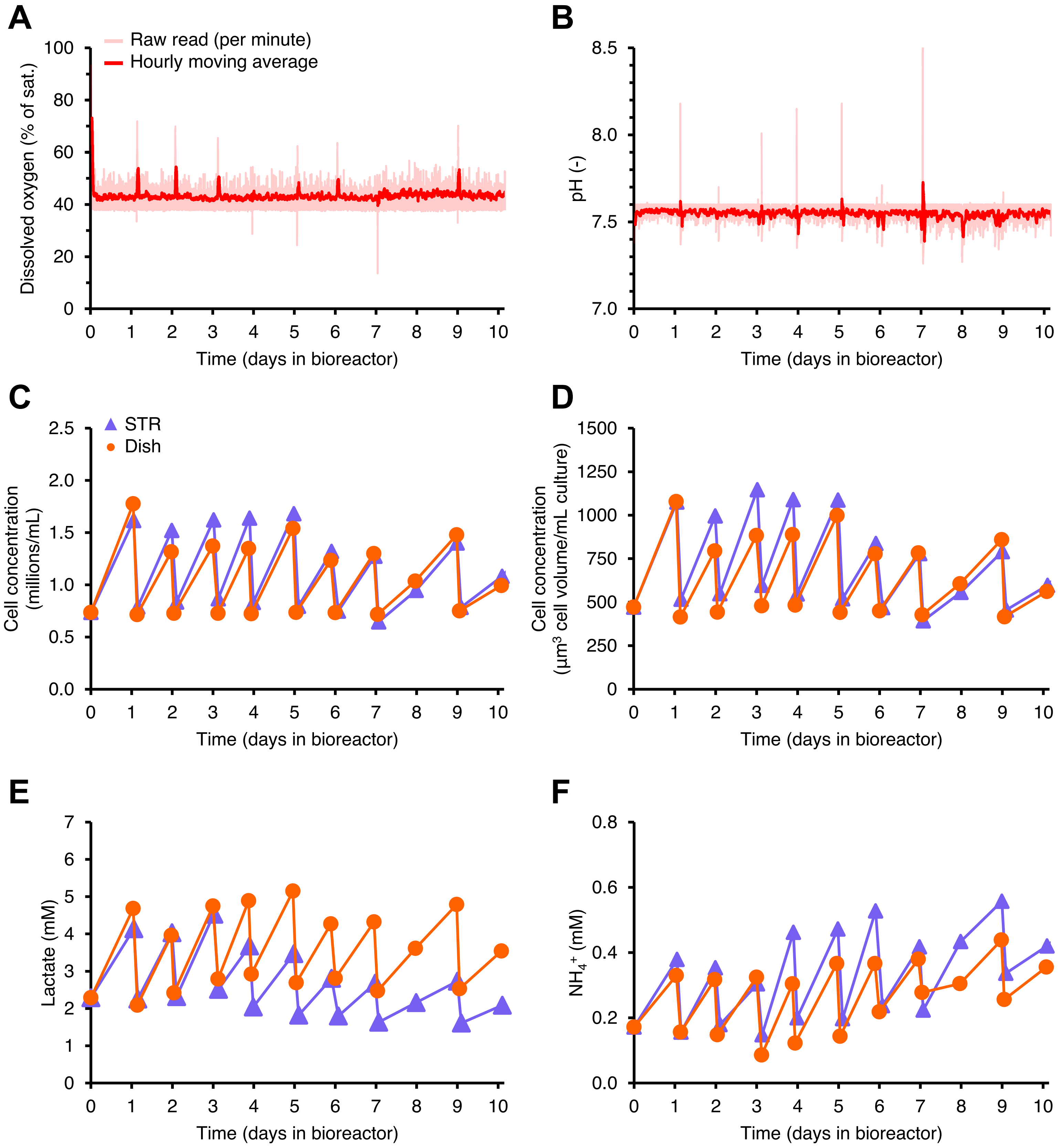


**Supplemental Figure S2. Expansion of erythroblasts in stirred tank bioreactors.** Erythroblasts were expanded from PBMCs for 9 days, and subsequently seeded in culture dishes or STRs (working volume = 300 mL; stirring speed = 200 rpm; marine down-pumping impeller with diameter = 2.8 cm; 100 mL/min $N_2$ headspace flow) at a starting cell concentration of $0.7\times10^6$ cells/mL. **(A)** $dO_2$ concentration was continuously measured and controlled at 40% (equivalent to 2.8 mg $O_2$/L) by sparging of air. **(B)** pH was continuously measured and controlled at 7.5 by sparging of $CO_2$. **(C)** Cell concentration was measured daily during 10 days of expansion. Cells were cultured following a sequential batch feeding strategy in which medium was refreshed if the daily measured cell concentration was $>1.2\times10^6$ cells/mL. **(D)** Total biomass concentration was calculated using the total cell volume (cells with diameter >5 µm) per mL of culture. **(E-F)** Extracellular lactate and ammonia concentrations were measured daily before and after medium refreshment. All data is displayed for a representative reactor run, using the same donor for both bioreactor and dish cultures.

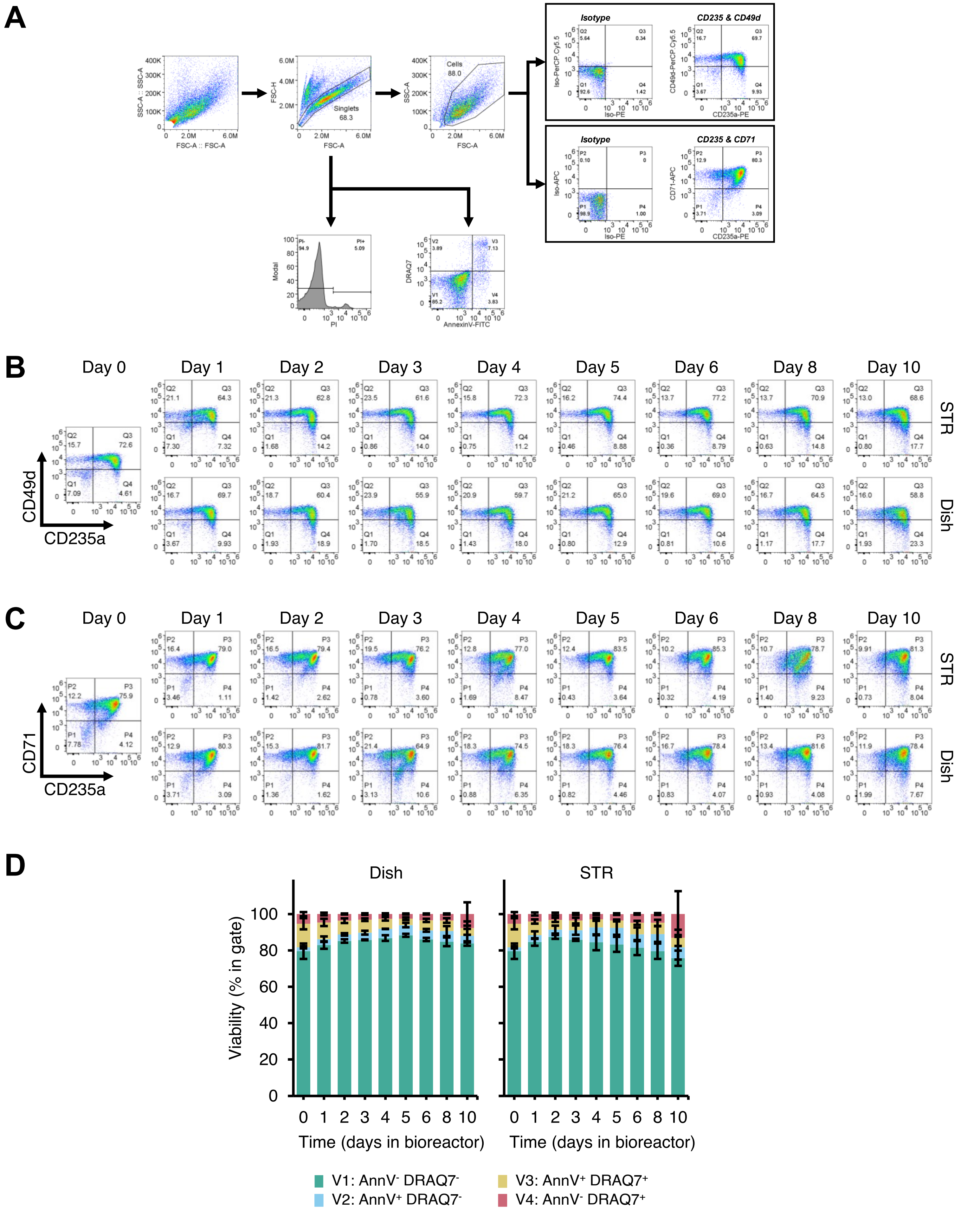


**Supplemental Figure S3. Erythroid cell surface marker expression and viability in erythroblasts expanded in stirred tank bioreactors.** Erythroblasts were expanded from PBMCs for 9 days, and subsequently seeded in culture dishes or STRs (working volume = 300 mL; stirring speed = 200 rpm; marine down-pumping impeller with diameter = 2.8 cm; 100 mL/min $N_2$ headspace flow) at a starting cell concentration of $0.7\times10^6$ cells/mL. **(A)** Gating strategy to evaluate the differentiation level of cultured erythroblasts. Single events were gated (FSC-A vs. FSC-H), followed by gating of cells (FSC vs SSC). Cells were stained for CD235a/CD71 or CD235a/CD49d, with gates defined based on appropriate IgG isotype controls. **(B-C)** Representative density plots indicating the expression of the cell surface markers CD71, CD235 and CD49d after 10 days of culture. **(D)** Cells were also stained with AnnexinV (apoptosis staining) and DRAQ7 (cell impermeable DNA stain); data displayed as mean ± SD (error bars; n=3 reactor runs / donors).

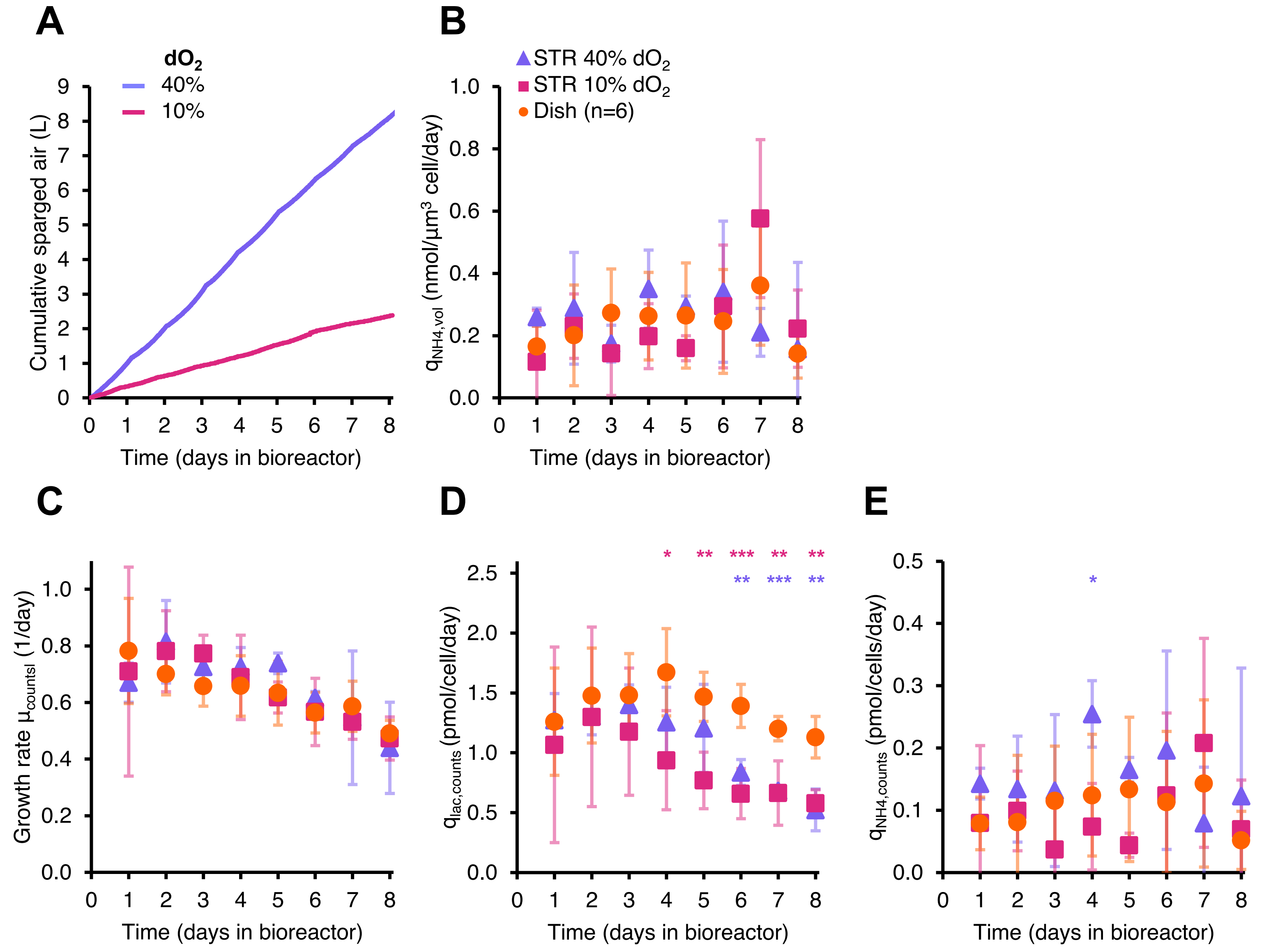


**Supplemental Figure S4. Lower dissolved oxygen setpoints lead to lower sparging requirements.** Erythroblasts were expanded from PBMCs for 9 days, and subsequently seeded in culture dishes or STRs (working volume = 300 mL; 200 rpm; pH=7.5; 100 mL/min $N_2$ headspace flow) at a starting cell concentration of $0.7\times10^6$ cells/mL. Dissolved oxygen was controlled by means of air sparging when below the targeted setpoint (10% of 40%, equivalent to 0.72 and 2.88 mg $O_2$/L respectively), and by stripping of excess oxygen using 100 mL/min $N_2$ headspace flow. **(A)** Cumulative volume of air sparged through the culture for representative reactor runs. **(B)** Cell-specific ammonium production rate ($q_{NH4,vol}$) calculated using growth rate data and measured extracellular lactate concentrations before and after each medium refreshment (see Supplemental Methods). **(C-E)** Rates were also calculated using the total cell concentration (cells per mL of culture). All data in panels **B-E** is displayed as mean ± SD (error bars; n=3 reactor runs / donors, unless indicated otherwise). Significance is shown for the comparison with dish cultures (unpaired two-tailed two-sample equal-variance Student's *t*-test; *$P<0.05$, **$P<0.01$, ***$P<0.001$, not displayed if difference is not significant).

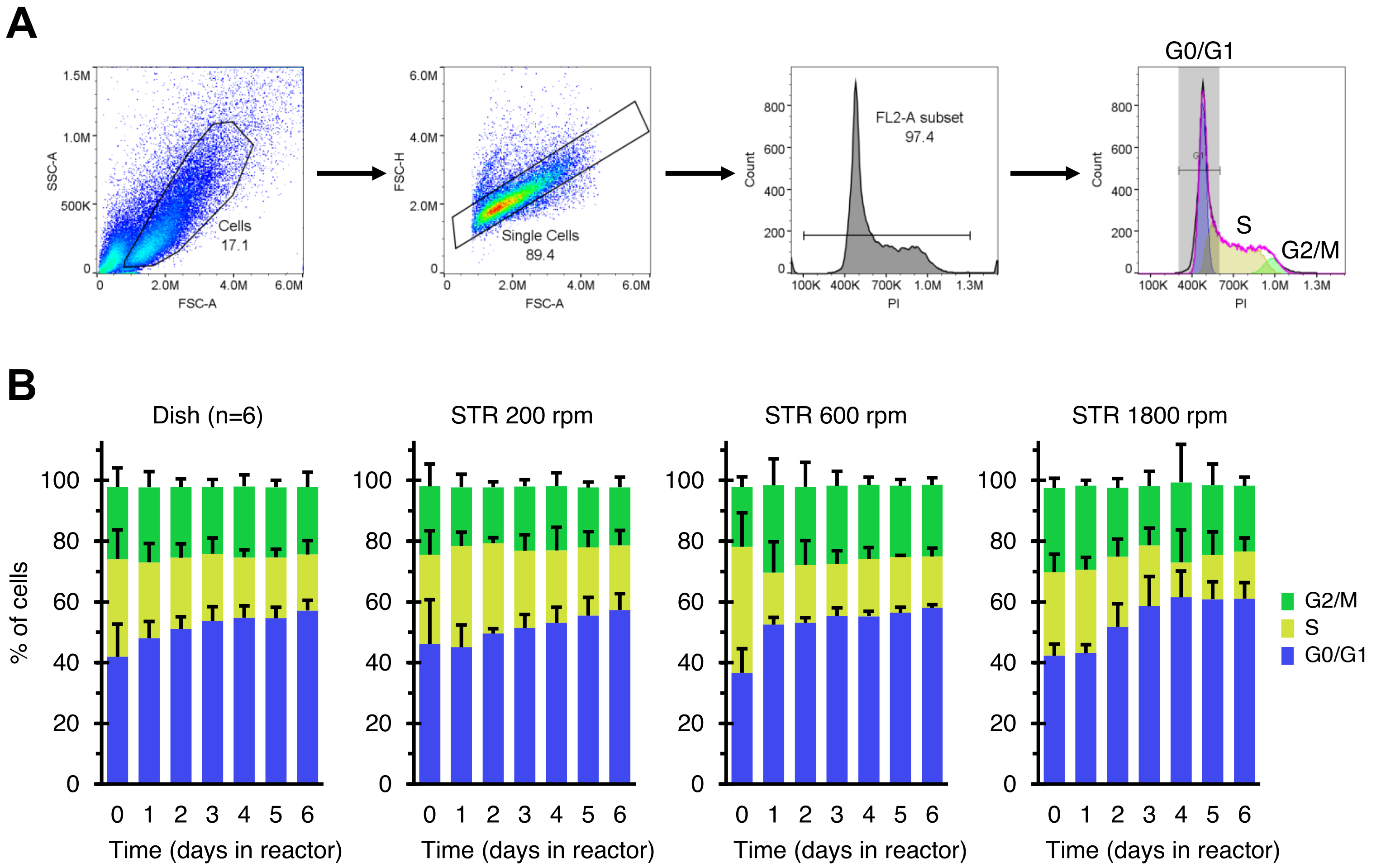


**Supplemental Figure S5. Effect of stirring speed on cell cycle of cultured erythroblasts.** Erythroblasts were expanded from PBMCs for 9 days, and subsequently seeded in culture dishes or STRs (working volume = 300 mL; marine down-pumping impeller with diameter = 2.8 cm; $dO_2$=40% controlled by sparging of air; pH=7.5 controlled by sparging of $CO_2$; 100 mL/min $N_2$ headspace flow) at a starting cell concentration of $0.7\times10^6$ cells/mL, under agitation at 200, 600 or 1800 rpm. **(A)** Gating strategy to evaluate the fraction of fixed cells in the G0/G1, S, or G2/M cell cycle stages. Cells were gated (FSC vs SSC), followed by gating of single events (FSC-A vs. FSC-H). Background in the PI channel was discarded, followed by fitting using the Watson Pragmatic algorithm. **(B)** Fraction of cells in the G0/G1, S, and G2/M cell cycle phases in bioreactor and static cultures. All data is displayed as mean ± SD (error bars; n=3 reactor runs / donors, unless indicated otherwise).

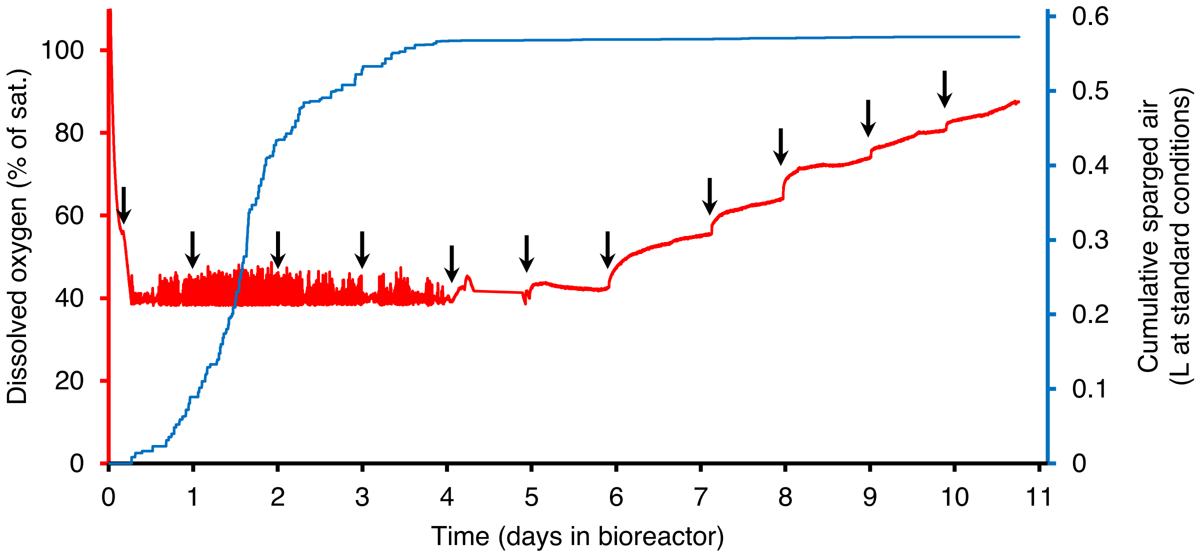


**Supplemental Figure S6. Oxygen requirements decrease during erythroblast differentiation.** Erythroblasts were expanded from PBMCs for 10 days, and subsequently seeded in differentiation medium at a starting cell concentration of $1 \times 10^6$ cells/mL. Cells were then transferred to culture dishes or STRs in culture dishes or STRs (working volume = 300 mL; stirring speed = 200 rpm; pH = 7.5) and kept in culture for 11 days without medium refreshment. Dissolved oxygen was measured continuously during the culture (red line; left axis). Air + 5% $CO_2$ was sparged only when $dO_2$ was <40%. Cumulative sparged air volume is also displayed (blue line; right axis). When taking samples (indicated with arrows), air was used to purge the sampling circuit, leading to an increase in the $dO_2$ concentration.

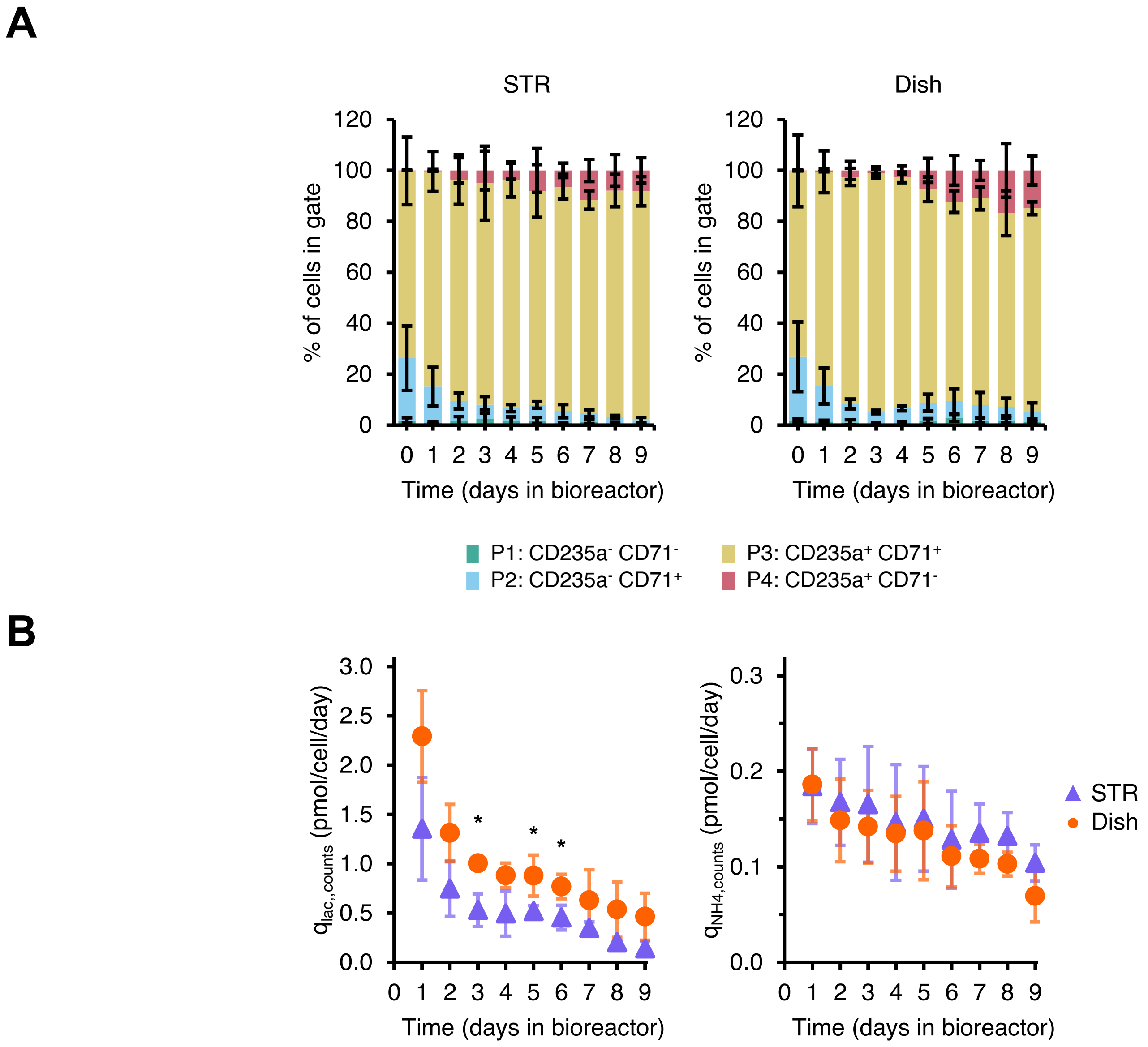


**Supplemental Figure S7. Expression of the erythroid surface markers CD235a and CD71 in erythroblasts cultured in 3 L stirred tank bioreactors.** Erythroblasts were expanded from PBMCs for 8 days, and subsequently seeded in culture dishes or 0.5 L stirred tank bioreactors at a starting cell concentration of 0.7-0.9×$10^6$ cells/mL. Cells were kept in culture following a fed batch feeding strategy in which medium was refreshed if the measured cell concentration (daily) was >1.2×$10^6$ cells/mL. Upon reaching a total number of >400 million cells, the culture was transferred to a 3.0 L bioreactor which was progressively filled by daily medium additions. **(A)** Cells were stained CD235a plus CD71 (erythroid differentiation markers). **(B)** Cell-specific production rates of lactate and ammonia calculated from the total cell counts, and extracellular metabolite concentrations. All data is displayed as mean ± SD (error bars; n=3 reactor runs / donors). Significance is shown for the comparison with dish cultures (unpaired two-tailed two-sample equal-variance Student's *t*-test; *$P$<0.05, **$P$<0.01, ***$P$<0.001, not displayed if difference is not significant).
