## Supplemental Materials for "Expansion and differentiation of *ex vivo* cultured erythroblasts in scalable stirred bioreactors"

### Supplemental methods

#### Cell cycle state measurements using flow cytometry

About $1\times10^6$ cells were centrifuged (5 minutes, 600g), washed with ice-cold PBS, fixed with 5 mL of ice-cold methanol, and stored at -20°C until analysis. From this suspension, 200.000 fixed cells were centrifuged (10 minutes, 1200g), washed with ice-cold PBS, resuspended in FxCycle™ PI/RNase staining solution (Invitrogen), incubated in the dark (10 minutes, 20°C), and analyzed using an Accuri C6 flow cytometer (BD Biosciences). Calculation of the percentages of cells in G0/G1, S, and G2/M phase was done using FlowJo™ using the Watson Pragmatic algorithm set to the following parameters: G1 peak range = 300.000 – 600.000; G1 peak CV = unconstrained; G2 peak = unconstrained; G2 peak CV = G1 peak CV.

#### Calculation of kinetic parameters

##### Growth rate and doubling time

For the expansion of erythroblasts, cells were cultured following a sequential batch feeding strategy in which medium was refreshed if the cell concentration (determined daily) was > $1.2\times10^6$ cells/mL. This leads to cell and metabolite concentrations over time as displayed in Supplemental Figure S2C-F. We assumed that cells followed exponential growth at a constant rate ($\mu_{max}$; 1/day) between consecutive dilution events:

$$\frac{d(C_N\,V_L)}{dt} = \mu_{max,counts}\,C_N\,V_L$$

where $C_N$ is the concentration of cells (in cells/mL), $V_L$ is the volume of culture (in mL), and $t$ is the time of culture (in days).

The maximum growth rate was estimated for each time interval between dilutions, using the measured cell concentration post- medium refreshment ($t_{i-1}$) and after overnight growth ($t_i$), assuming no change in culture volume between both timepoints (i.e. evaporation)::

$$\mu_{max,counts}(t_i) = \frac{\ln\left(\frac{C_N(t_i)}{C_N(t_{i-1})}\right)}{t_i - t_{i-1}}$$

As cell size decrease is characteristic of erythropoiesis, the total volume of cells per unit of volume of culture ($C_V$; μm$^3$ of cells per mL of culture) was also used as proxy for total biomass concentration. With this biomass concentration, we calculated a growth rate that takes into account the progressive decrease in cell size during culture ($\mu_{max,vol}$; 1/day):

$$\mu_{max,vol} = \frac{\ln\left(\frac{C_V(t_{i+1})}{C_V(t_i)}\right)}{t_{i+1} - t_i}$$

For both $\mu_{max,counts}$ and $\mu_{max,vol}$, only data of cells with a diameter larger than 7.5 or 5 μm was used, respectively. Doubling time was calculated as:

$$\tau = \frac{\ln 2}{\mu_{max}}$$

### Cell-specific consumption and production rates of metabolites

Cell-specific consumption/production rates of metabolites were assumed to be constant between consecutive medium refreshment events, and were calculated using metabolite concentration data and the calculated growth rate:

$$\frac{d(C_{lac}\, V_L)}{dt} = q_{lac,counts}\, C_N\, V_L$$

For instance, the cell-specific production rate of lactate ($q_{lac,counts}$; mol lactate/cell/day) was calculated as:

$$q_{lac,counts}(t_i) = \mu_{max,counts}(t_i)\, \frac{C_N(t_i) - C_N(t_{i-1})}{C_{lac}(t_i) - C_{lac}(t_{i-1})}$$

where $C_{lac}(t_i)$ is the measured supernatant concentration of lactate (in mol/mL) at timepoint $t_i$. Rates were also calculated using the total biomass concentration (e.g. $q_{lac,vol}$; mol lactate/ μm$^3$ cell/day).

### Cell-specific oxygen consumption rate

For the determination of the cell-specific oxygen consumption rate ($q_{O2}$; mol $O_2$/cell/h), dissolved oxygen data between consecutive medium refreshments (~24 h intervals) was fitted to a black box model assuming a constant growth rate and $q_{O2}$ in the analyzed time interval:

$$\frac{d(c_{O2}\, V_L)}{dt} = k_L a(c_{O2}^{sat} - c_{O2})V_L - q_{O2} C_N V_L$$

where $V_L$ is the culture volume (in L), $c_{O2}^{sat}$ is the saturation oxygen concentration for the culture conditions (7.2 mg $O_2$/L = 0.225 mmol $O_2$/L using air + 5% $CO_2$ at atmospheric pressure and 37°C), $c_{O2}$ is the dissolved oxygen concentration in the culture (in mol/L, calculated as $c_{O2} = d_{O2} \times c_{O2}^{sat}$), and $k_L a$ is the overall volumetric mass transfer coefficient of the system recalculated to a liquid-side mass transfer oxygen composition gradient (with $a$ being the specific interfacial area = $m^2$ interfacial area/$m^3$ liquid volume) (Shuler & Kargi, 2002). The cell concentration $C_N$ (cells/L) was calculated for all timepoints of each day using the estimated growth rate:

$$C_N(t) = C_{N,0}\, e^{\mu_{max,count}\, t}$$

The value of $q_{O2}$ for each day was calculated by iteratively solving the differential equation describing $c_{O2}$ was and fitting to each 24 h interval of $d_{O2}$ measurements. The value of the overall mass transfer coefficient of the system ($k_L a$) was determined experimentally for the culture conditions (using water, with equal working volume, temperature and pressure) with the dynamic method: oxygen was purged from the liquid by nitrogen sparging until reaching a $dO_2$ <0.5%, followed by a constant flow of air in the headspace of 100 mL/min until close to saturation ($dO_2$>90%). The value of $k_L a$ was estimated by fitting of the same black box model to the measured $dO_2$ data, with $q_{O2} = 0$ (Bandyopadhyay et al., 1967; Tribe et al., 1995).
